# Oncogenic alterations in the p53 pathway abolish oscillatory competence

**DOI:** 10.1101/2021.02.09.430245

**Authors:** Lingyun Xiong, Alan Garfinkel

## Abstract

The tumor suppressor p53 displays concentration oscillations in response to DNA damage, a behavior that has been suggested to be essential to its anti-cancer function. Many genetic alterations in the p53 pathway have been shown to be oncogenic, whether by experiment or by clinical associations with various cancers. These oncogenic alterations include somatic mutations, copy number variations and inherited polymorphisms. Using a differential equation model of p53-Mdm2 dynamics, we employ Hopf bifurcation analysis to show that all of the oncogenic perturbations have a common effect, to abolish the oscillatory competence of p53, thereby impairing its tumor suppressor function. In this analysis, these diverse genetic alterations, widely observed in human cancers, have a unified mechanistic explanation.

**Significance:** In human cancers, the p53 tumor suppressor pathway is frequently altered by diverse genetic changes. An integrated understanding of these oncogenic alterations is currently lacking. We show that all oncogenic alterations in the p53 pathway abolish the oscillatory competence of p53, a property that is essential for cell cycle arrest upon stress, for effective DNA damage response and for maintaining genome integrity. This unified dynamical explanation of distinct cancer driver events that converge on a key cancer hallmark pathway has practical implications for anti-cancer therapies.

## 1 Introduction

The tumor suppressor p53 plays important roles in safeguarding the genome against genotoxic and other stresses, thereby preventing cancer (P. Chen, Chen, Bookstein, & Lee, 1990; Finlay, Hinds, & Levine, 1989; A. J. Levine, 1997). p53 has several inhibitors, including Mdm2 (Haupt, Maya, Kazaz, & Oren, 1997; Momand, Zambetti, Olson, George, & Levine, 1992). Oren, Levine and colleagues showed that a simple model of p53-Mdm2 negative feedback predicted oscillations in p53 protein concentration, which they confirmed by observing one cycle of the p53-Mdm2 oscillation after ionizing radiation in cultured cells (Lev Bar-Or et al., 2000). Lahav and colleagues found that 7-irradiation triggered multiple oscillations in p53 and Mdm2 in single cells, with a period of about 5.5 hours (Lahav et al., 2004); they also developed an explanatory mathematical model (Geva-Zatorsky et al., 2006).

Oscillations in the p53-Mdm2 system have been suggested to have functional roles, as a mode of intracellular communication to coordinate cell cycle arrest and DNA repair, which crucially determine cell fate after DNA damage (Lev Bar-Or et al., 2000; Purvis et al., 2012). As a result, impaired p53 oscillations would compromise genome integrity and promote cancer.

Genetic alterations in the p53 pathway are frequently oncogenic, including somatic mutations, copy number variations and inherited polymorphisms affecting p53 and its regulators such as Mdm2, Mdm4 and ATM. Here, we show that each of these oncogenic genetic changes, when biophysically modeled in the p53-Mdm2 negative feedback system, acts to abolish the oscillations upon DNA damage, thus providing an insight into the common mechanism of their oncogenic effect: a loss of the oscillatory competence of p53.

## 2 Results

### The mathematical model

To study the effect of individual oncogenic alterations on the oscillatory behavior of the p53-Mdm2 system, we developed a mathematical model of the core p53-Mdm2 negative feedback system. Based on previous studies (Batchelor, Mock, Bhan, Loewer, & Lahav, 2008; Geva-Zatorsky et al., 2006; Heltberg et al., 2019), our model reflects the transcription factor p53 enhancing the production of its own inhibitor Mdm2 (Fig. 1A):

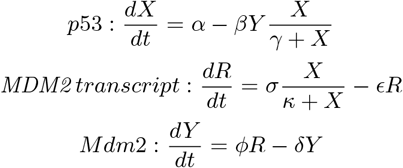

**Figure 1:**
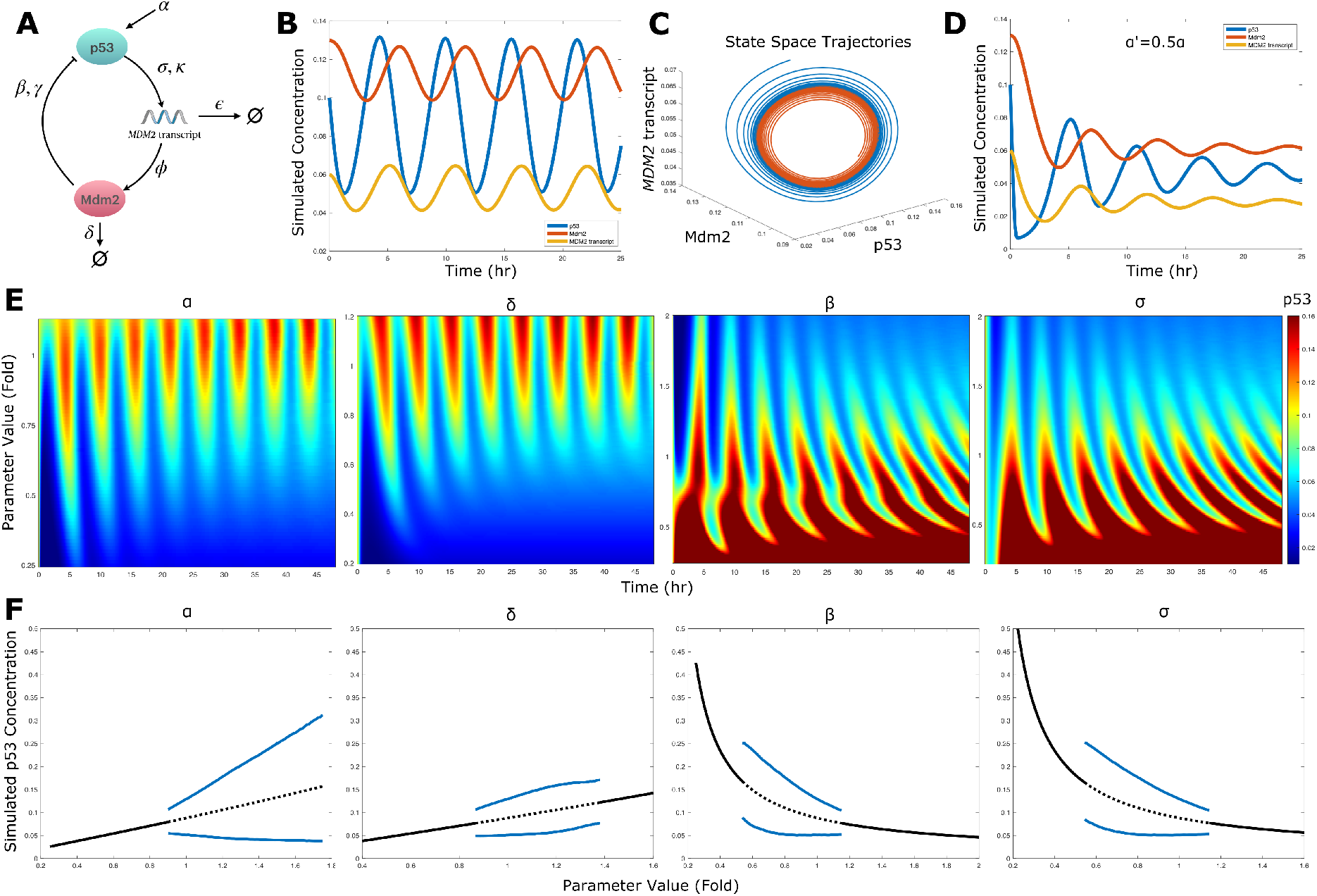
The oscillatory behavior of the p53-Mdm2 system during DNA damage response is sensitive to parameter values. (A) Schematic of the proposed model. (B) Numerical simulation of the model in response to DNA damage, showing simulated p53 protein level (blue), *MDM2* transcript level (yellow), and Mdm2 protein level (red). (C) The long-term behavior of the system approaches a limit cycle attractor that is stable under perturbations (the blue and red trajectories). (D) p53 pulses in response to DNA damage are dampened when *α* is halved. (E) Simulated p53 concentrations over time (the horizontal axes) are shown as heatmaps for varying parameter values of *α, δ*, *β* and *σ*. (F) One-dimensional bifurcation diagrams of the p53 system against single-parameter variation in *α, δ, β* and *σ*. The steady-state p53 concentration is shown as a solid line when it is stable and dotted when unstable; the amplitudes of the limit cycle oscillations are indicated by the blue curves.

The key features of the mathematical model are: (i) p53 is produced at a constant rate (*α*) and is degraded upon binding to Mdm2 through a saturated degradation process (*β, γ*), where *β* represents the maximum rate of Mdm2-induced degradation of p53 and *γ* the p53 concentration for half-maximum degradation; (ii) *MDM2* transcript is produced upon p53 transactivation through a saturated induction process (*σ, κ*) and degraded through a first-order decay process (*ϵ*), where *σ* represents the maximum rate of p53-dependent transcription of *MDM2* and *κ* the p53 concentration for half-maximum *MDM2* transcription; (iii) Mdm2 is produced proportional to the *MDM2* transcript level (*ϕ*) and degraded through a first-order decay process (*δ*).

We chose appropriate values for each biophysical parameter based on previous calibrations *(Methods*). Numerical simulation of the mathematical model recapitulated the oscillatory behavior of p53 during DNA damage response, and agreed with experimental observations that p53 protein level oscillates with a characteristic period of 5-6 hours, and that both *MDM2* transcript and Mdm2 protein levels oscillate during DNA damage (Fig. 1B-C) (Batchelor et al., 2008; Geva-Zatorsky et al., 2006; Hanson, Porter, & Batchelor, 2019; Heltberg et al., 2019; Lahav et al., 2004; Purvis et al., 2012).

We evaluated known oncogenic alterations in the p53 pathway for their effect on the biophysical parameters of the p53-Mdm2 system (Table 1). They affect three essential aspects of p53-Mdm2 dynamics: production of p53 (*α*), p53-Mdm2 interactions (*β* and *σ*), and Mdm2 degradation (*δ*).

**Table 1:**
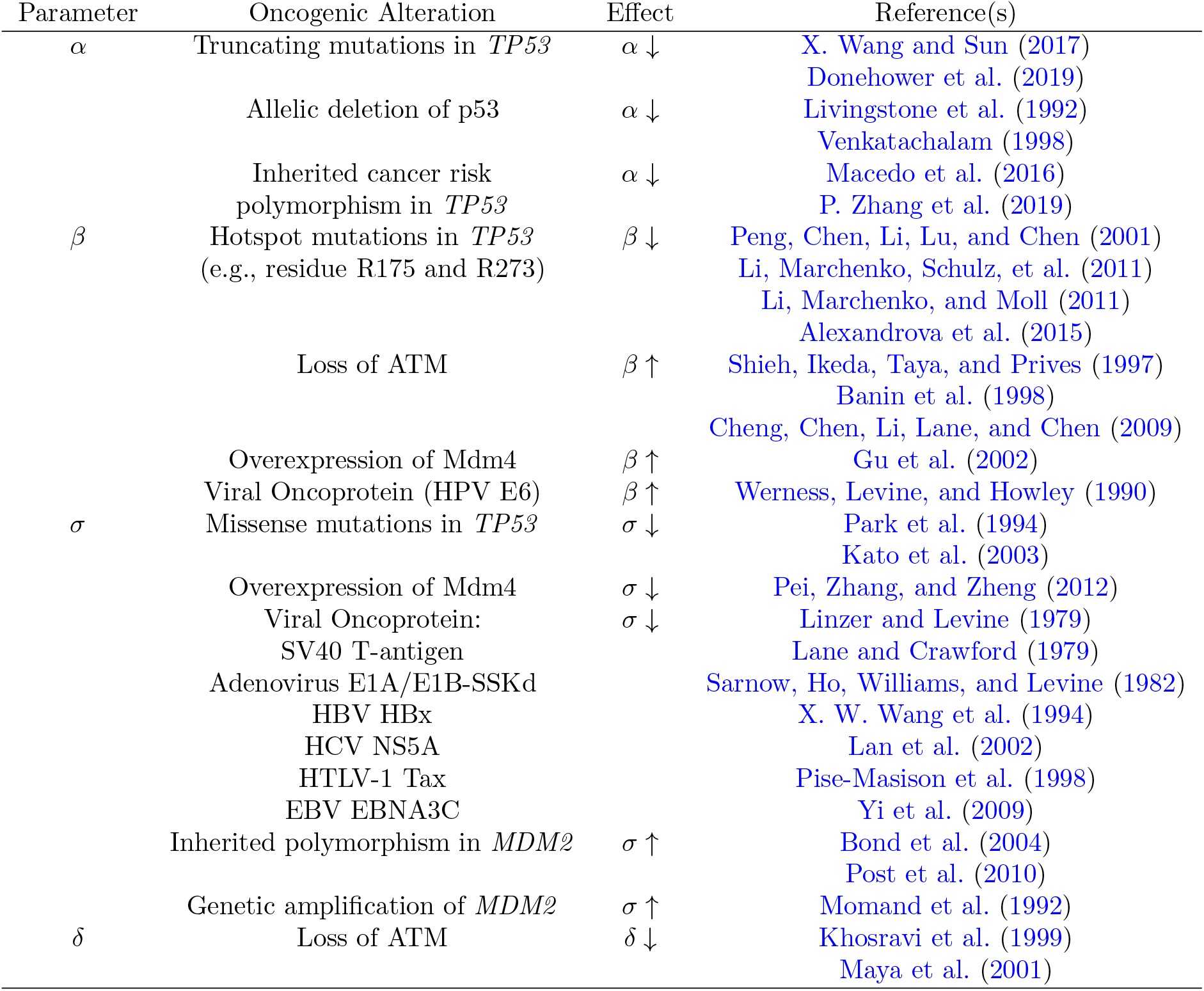
Impact of oncogenic alterations on model parameters.

### The oscillatory behavior of p53 is sensitive to parameter values

We studied the transient response of the p53-Mdm2 system by performing a series of single-parameter sensitivity analyses for *α, β, σ* and *δ* (*Methods*). We found that lowering *α* reduced the amplitude of initial p53 pulses in response to DNA damage (Fig. 1E); a 50% reduction of *α* resulted in markedly reduced initial amplitudes and damped oscillations in both p53 and Mdm2 (Fig. 1D). Decreasing *δ* had a similar effect (Fig. 1E). Lowering *β* or *σ* dramatically enhanced initial p53 pulses and shortened their latency. Further lowering of *β* and *σ* increased the oscillatory period and ultimately caused a loss of the oscillatory response. Increasing *β* or *σ*, on the other hand, reduced initial p53 pulses and diminished the oscillatory behavior of the system (Fig. 1E).

The long-term behavior of the system displayed sustained p53 oscillations only when *α* was sufficiently large (Fig. 1F). For *β, σ* and *δ*, however, we noted that robust p53 oscillations first emerged and then disappeared as parameter values varied (Fig. 1F), consistent with previous numerical studies on related models (Ciliberto, Novak, & Tyson, 2005; Lev Bar-Or et al., 2000; Ma et al., 2005; Wagner et al., 2005). That is, p53 oscillations were viable only within a range of parameter values, a “Goldilocks” effect that is explained by these results. Each one of these biophysical parameters can affect the oscillatory competence of the p53-Mdm2 system.

### Parametric conditions for sustained p53 oscillations

Next, we sought to understand how the system would respond to simultaneous changes in multiple parameters. A single oncogenic event can alter more than one biophysical parameter, such as p53 hotspot mutations and the loss of ATM (Table 1). Also, multiple oncogenic events can co-occur in tumors, for instance, p53 mutations and loss of heterozygosity on chromosome 17p (Donehower et al., 2019).

Moreover, while simulations can reveal the presence of periodicity, it is important to ask whether a true qualitative change has occurred, leading to the presence of a *limit cycle attractor,* with a preferred frequency and, importantly, stable under a variety of perturbations and changes of initial conditions. These are all highly desirable properties for the cell’s DNA damage response system (for this point see also Wagner et al. (2005).

We therefore looked for limit cycle attractors in our model using Hopf Bifurcation Theory (*Methods*). There is only one equilibrium point of the p53-Mdm2 system at which p53, *MDM2* transcript and Mdm2 levels are all positive. By evaluating the eigenvalues of the Jacobian matrix at this equilibrium point, we derived the criteria for the emergence of a limit cycle attractor:

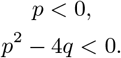

Here, –*p*/2 is the real part of the pair of complex conjugate eigenvalues; *p*^2^ – 4*q* < 0 is required for the eigenvalues to be complex conjugate; *p* and *q* are functions of model parameters.

Note that p53 oscillations emerge by subcritical bifurcations (Fig. 1F), meaning that they emerge full-blown at a finite amplitude, rather than growing slowly from zero. This is also desirable for the DNA damage response system, as was pointed out by Ciliberto et al. (2005). Two types of bifurcation have this property: subcritical Hopf bifurcation and saddle-node-loop bifurcation. Saddle-node-loop bifurcation is ruled out here because there is only one equilibrium point at all parameter value combinations we examined.

### Loss of ATM jeopardizes p53 oscillations upon DNA damage

We used the criteria for the emergence of a limit cycle attractor to study how ATM regulates the p53-Mdm2 system in response to DNA damage. When cells are non-stressed, p53 concentration is maintained at very low levels. Following DNA double strand breaks (DSBs), p53 and Mdm2 exhibit oscillatory behavior, which can last for several days (Batchelor et al., 2008; Geva-Zatorsky et al., 2006). Upon DSBs, ATM activates p53 by inhibiting Mdm2-mediated p53 degradation (primarily lowering *β* (Banin et al., 1998; Cheng et al., 2009; Shieh et al., 1997)) and mediating fast degradation of Mdm2 (increasing *δ* (Khosravi et al., 1999; Maya et al., 2001; Stommel & Wahl, 2004)) (Fig. 2A). To examine the joint effect of these two biophysical processes, we performed Hopf bifurcation analysis using *β* and *δ* as the bifurcation parameters, while keeping other parameters constant. Intriguingly, the conditions for Hopf bifurcation are satisfied within an ‘island’ in *β*-*δ* parameter space (Fig. 2C; Fig. S1A, B).

**Figure 2:**
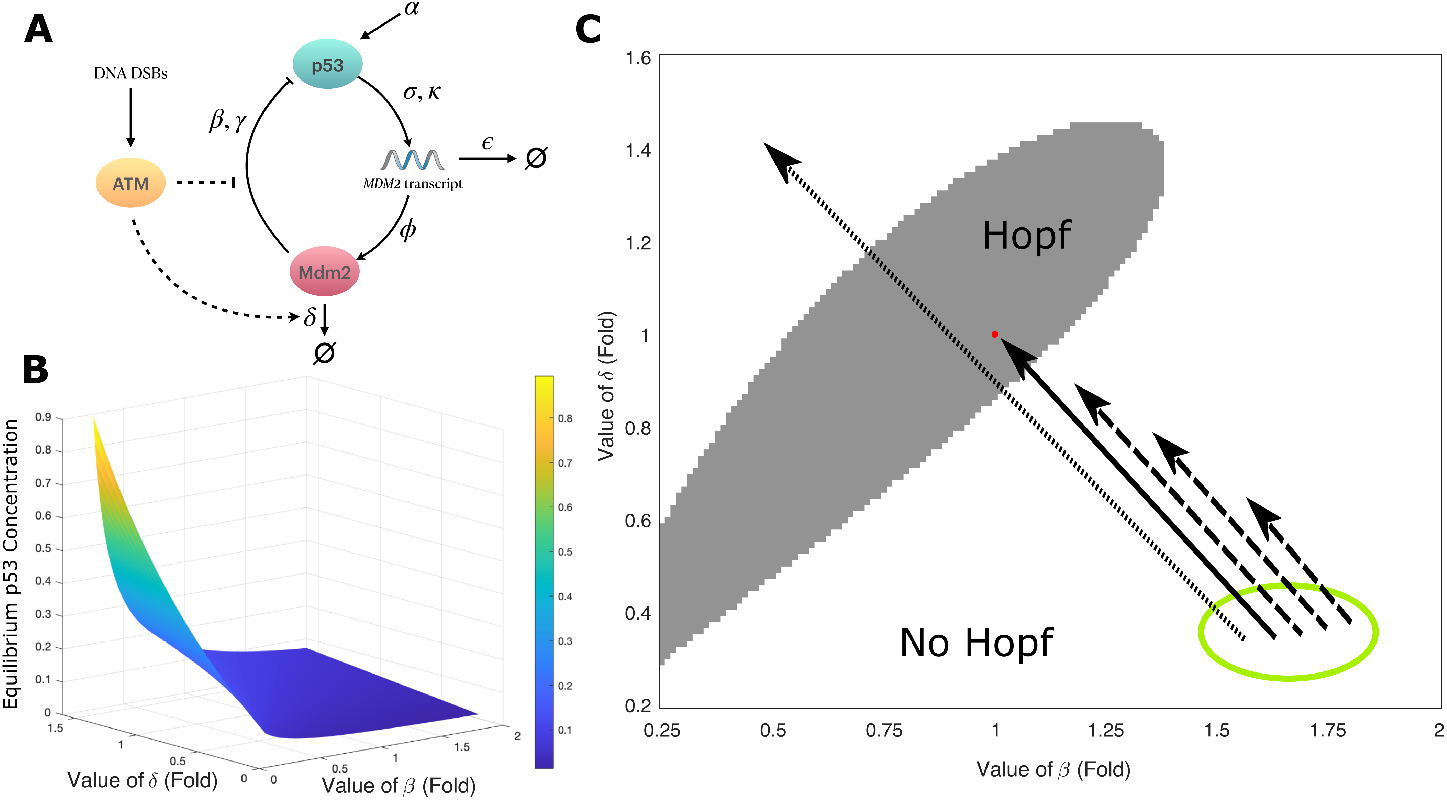
ATM mediates the onset of p53 oscillations. (A) The impact of ATM on the p53-Mdm2 system in response to DNA DSBs. (B) The equilibrium concentration of p53 as a function of parameters *β* and *δ*, describing a surface over the *β*-*δ* plane. (C) Hopf bifurcation analysis identifies the critical region in the *β*-*δ* parameter space, where the p53-Mdm2 system displays a stable oscillatory response to DNA damage. The gray region specifies the parameter subspace where p53 oscillations are self-sustained (*p* < 0 and *p*^2^ — 4*q* < 0); while p53 concentration goes to equilibrium in the white region. The green circle denotes a hypothetical region where the p53-Mdm2 system operates under non-stressed conditions. The red dot denotes the best-fit combination of *β* and *δ* for DSB conditions. ATM activates p53 by shifting the state of the system into the critical region (the solid arrow). Loss of ATM would compromise this transition (dashed arrows); hyperactive ATM results in over-transition (the dotted arrow). DSBs: double strand breaks.

We therefore suggest that the action of ATM in creating the oscillatory response upon DSBs can be explained as a Hopf bifurcation induced by ATM’s impact on lowering *β* (Mdm2 inhibition of p53) and increasing *δ* (degradation of Mdm2), representing a shift of system state in *β*-*δ* parameter space (the solid arrow in Fig. 2C), from a non-oscillatory region (the green circle in Fig. 2C) to the critical region where oscillation is self-sustained (the gray region in Fig. 2C). The best-fit parameter combination (the red dot in Fig. 2C) resides inside, but near the boundary of the critical region. It is plausible that reduced levels of ATM, due to mutations or copy number loss, is insufficient to carry the system into the critical region after DNA damage (dashed arrows in Fig. 2C), rendering the p53-Mdm2 system incapable of oscillation. And indeed, ATM inhibitors dampen p53 oscillation upon treatment with the radiomimetic drug neocarzinostatin in multiple cancer cell lines with wild type p53 (Stewart-Ornstein & Lahav, 2017). On the contrary, DNA-dependent protein kinase inhibitors that *hyper*activate ATM increase p53 abundances but do not trigger oscillations (Stewart-Ornstein & Lahav, 2017), possibly due to over-transiting the system beyond the critical region (the dotted arrow in Fig. 2C; Fig. 2B).

In contrast to the dual effect of ATM on model parameters, a horizontal shift leftward from the green region in the parameter space alone would bring the system closer to the Hopf bifurcation zone, to a region where the system has a damped oscillatory response (Fig. 2C). It would also result in an increase in p53 levels (Fig 2B). These two changes combine to produce a prolonged pulse of high amplitude (see Fig 1E for low values of *β*), as seen in ATR-mediated p53 response to ultraviolet radiation (Batchelor, Loewer, Mock, & Lahav, 2011).

### Oncogenic alterations abolish p53 oscillations

The remaining oncogenic alterations affect the parameters *α, β* and *σ* (Table 1). Considering that *β* and *σ* co-occur as a product throughout the analytic calculation (*Methods*), we let *α* and *βσ* be the bifurcation parameters for the Hopf bifurcation analysis while keeping the other parameters constant. The equilibrium point of this system (Fig. 3A) was stable when the values of *α* or *βσ* were relatively small. Importantly, we identified an area in the *α*-*βσ* plane where the parametric conditions of Hopf bifurcation were fulfilled, ensuring robust p53 oscillation (Fig. S2). Similarly, Hopf bifurcation analysis on *α*, *βσ* and *δ* revealed a Hopf subspace (Fig. 3C). Note that the best-fit parameter combination (the red sphere) is located within the Hopf subspace, but fairly near the boundary, where a modest shift in parameter values could result in a qualitative change in the dynamic behavior of the p53-Mdm2 system, resulting in a loss of p53 oscillations.

**Figure 3:**
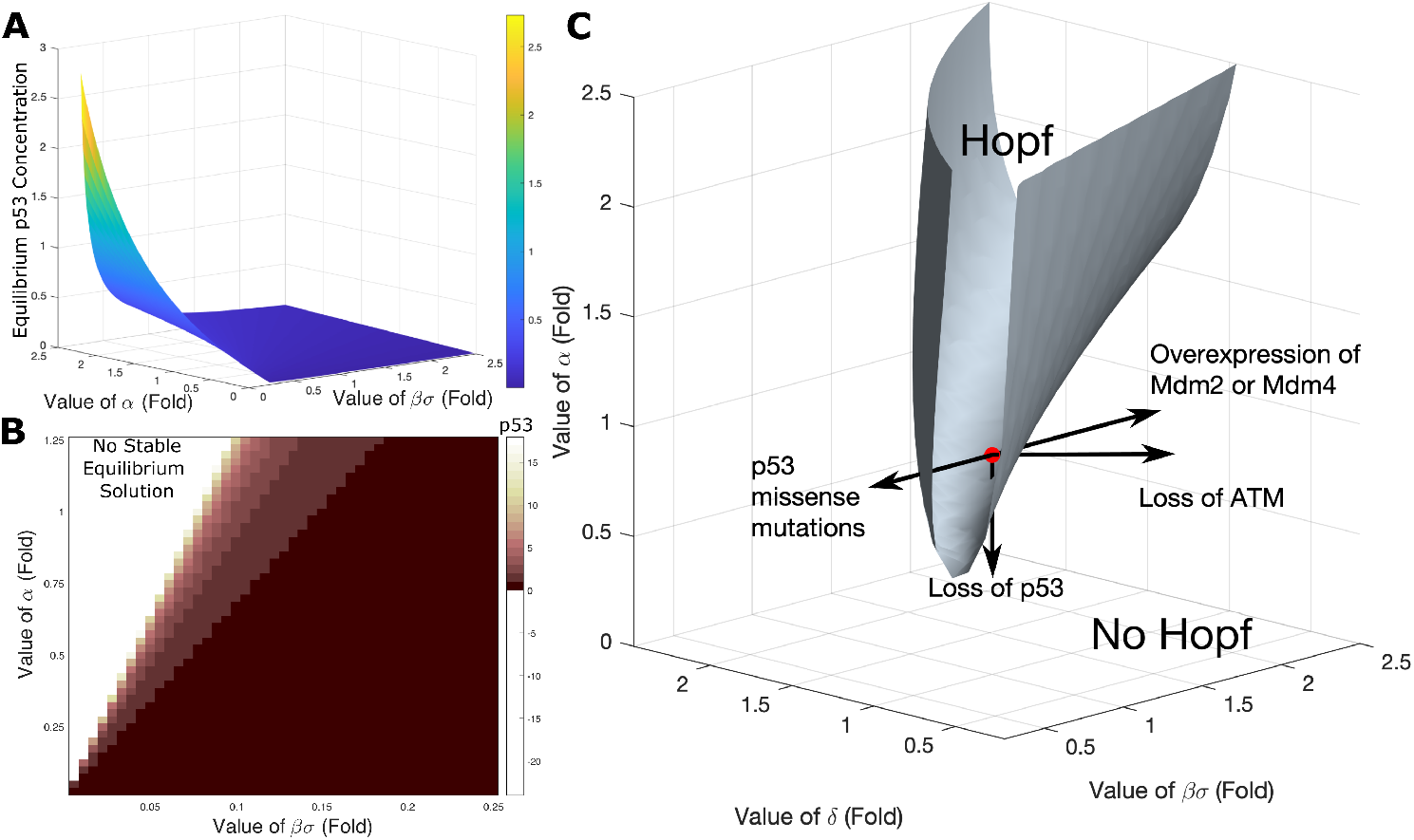
Oncogenic alterations take multiple routes to abolish p53 oscillations. (A) The equilibrium concentration of p53 is a function of parameter *α* and *βσ*, which describes a surface over the *α-βσ* plane. (B) For small values of *βσ*, the equilibrium concentrations of p53 are shown in a heatmap. The p53-Mdm2 system loses its stable equilibrium point when *βσ* is extremely small (white region), and the p53 concentration grows unbounded in time during DNA damage. (C) Hopf bifurcation analysis, by calculation of eigenvalues, indicates that oscillatory behavior of the p53-Mdm2 system is self-sustained in a parameter subspace of *α, βσ* and *δ*, enclosed by the gray surface and annotated as ”Hopf” (*p* < 0 and *p*^2^ – 4*q* < 0). The red sphere marks the best-fit combination of parameter values. The arrows indicate the impact of broad-category oncogenic alterations on parameter values.

In this dynamical systems framework, there are four routes for diverse oncogenic alterations to impair p53 oscillations during DNA damage response (Fig. 3C; Movie S1).

First, p53 missense mutations reduce the p53-dependent *MDM2* transcription rate (lowering *σ* (Kato et al., 2003; Park et al., 1994); p53 hotspot mutations also reduce the Mdm2-dependent p53 degradation rate (lowering *β* (Alexandrova et al., 2015; Li, Marchenko, & Moll, 2011; Li, Marchenko, Schulz, et al., 2011; Peng et al., 2001)), pushing the system out of the Hopf subspace (Fig. 3C). We observed that small values of *βσ* result in high steady-state p53 levels (Fig. 3A), which explains the accumulation of mutant p53 in tumor-derived cells as well as the elevated p53 levels in cells transformed by tumor viruses such as SV40 and adenovirus type 5 (Table 1). For extremely small values of *βσ*, p53 levels grow without bound (Fig. 3B), reconciling the observation that the Mdm2 inhibitor Nutlin-3, known to block Mdm2-dependent p53 degradation, potently abrogates p53 oscillations and increases p53 levels greatly (Purvis et al., 2012; Stewart-Ornstein & Lahav, 2017). In addition, given the span of the Hopf subspace for lower *βσ* values (Fig. S2; Movie S1), we reason that p53 missense mutations that moderately reduce the transactivation activity of p53 would be tolerated and have limited pathological consequences, as has been reported in functional screens of p53 mutants (Giacomelli et al., 2018; Kotler et al., 2018). In these cases, an additional loss of p53 would assist to compromise the oscillatory competence of the p53-Mdm2 system, plausibly accounting for the prevalent loss of heterozygosity in tumors with mutant p53 (Donehower et al., 2019).

Second, a loss of p53, due to inherited cancer risk polymorphisms, truncating mutations or allelic deletion in *TP53,* decreases the production rate of p53 (lowering *α* (Donehower et al., 2019; Livingstone et al., 1992; Macedo et al., 2016; Venkatachalam, 1998; X. Wang & Sun, 2017; P. Zhang et al., 2019)), dropping the system out of the Hopf subspace (Fig. 3C). Phenotypically, the loss of p53 oscillations due to germline mutations in *TP53* could account for the contribution of genotoxic therapies to the development of subsequent primary tumors among individuals with Li-Fraumeni syndrome (Bougeard et al., 2015; Kasper et al., 2018).

Third, genetic amplification or a certain inherited polymorphism increases *MDM2* transcription rate (elevating *σ* (Bond et al., 2004; Momand et al., 1992; Post et al., 2010)), readily disrupting p53 oscillations upon DNA damage (Hu et al., 2007). Overexpression of Mdm4 due to inherited polymorphisms or copy number gain has a net effect of increasing *βσ* (Table 1; *SI Appendix*). Of note, the viral oncoprotein HPV E6 enhances p53 degradation (increasing *β* (Scheffner, Werness, Huibregtse, Levine, & Howley, 1990; Werness et al., 1990)). These alterations all bring the system out of the Hopf subspace (Fig. 3C).

Lastly, we showed that ATM mediates the onset of p53 oscillations through its impact on β and δ (Fig. 2C), thus a loss of ATM keeps the system out of the Hopf subspace during DNA damage response (Fig. 3C).

*Together, despite that oncogenic alterations take multiple routes to alter key biophysical parameters of the p53-Mdm2 system, they have the common effect: p53 oscillations are abolished.*

## 3 Discussion

Physiological processes were long thought to be governed by static equilibria. But that concept, the doctrine of homeostasis, is giving way to a more dynamical view of physiological processes, many of which are characterized by oscillatory modes of operation (Boiteux, Hess, & Sel’Kov, 1980; Ferrell, Tsai, & Yang, 2011; J. H. Levine, Lin, & Elowitz, 2013). The emerging picture is that health consists of the successful coordination of many oscillatory processes; conversely, disease may often manifest as a loss of these oscillatory processes and/or a loss of their coordination (Buhr, Yoo, & Takahashi, 2010; Glass, 2001; Noble, 2006; Tu, Kudlicki, Rowicka, & McKnight, 2005).

Our results, together with previous findings, suggest that this may well be the case for p53, since, in our model, the functional response to DNA damage is to generate oscillations in the p53-Mdm2 system, whereas oncogenic alterations all disrupt p53-Mdm2 oscillations, as predicted by Hopf bifurcation theory. Thus, genetic alterations related to cancer, which were largely established through correlation or experimental manipulation, can now be understood mechanistically: diverse oncogenic alterations in the p53 pathway drive cancer by abolishing p53-Mdm2 oscillations.

### Hypothesis: A Role For ARF

Besides coordinating DNA damage response, p53’s role as a tumor suppressor also manifests in response to oncogenic signaling such as the overexpression of Myc, E1A, or E2F-1 (D. Chen, Shan, Zhu, Qin, & Gu, 2010; Efeyan, Garcia-Cao, Herranz, Velasco-Miguel, & Serrano, 2006). Upon oncogenic activation, the tumor suppressor ARF triggers p53-dependent cell cycle arrest and/or apoptosis (Kamijo et al., 1997). Mice lacking ARF are highly prone to tumor development (Kamijo, Bodner, van de Kamp, Randle, & Sherr, 1999), and deficiency in ARF underlies multiple cancer types in humans (Knijnenburg et al., 2018; Sherr, 1998). ARF activates p53 by inhibiting Mdm2-mediated degradation of p53 and accelerating Mdm2 degradation (Pomerantz et al., 1998; Weber, Taylor, Roussel, Sherr, & Bar-Sagi, 1999; Y. Zhang, Xiong, & Yarbrough, 1998). Since this is also the mechanism of ATM-induced p53 oscillations during the DNA damage response (Fig. 2), it is likely that ARF triggers p53 oscillations in response to oncogenic activation. If true, a loss of ARF could also abolish the oscillatory competence of the p53-Mdm2 system, thus promoting oncogenesis.

### Therapeutic Implications

Small molecules APR-246 and COTI-2 can reactivate missense-mutant p53 by reversing its oncogenic effect of lowering *σ*, thereby bringing the p53-Mdm2 system back to the oscillatory regime (Bykov et al., 2002; Lambert et al., 2009; Salim, Maleki Vareki, Danter, & Koropatnick, 2016). Their therapeutic efficacy is currently being tested in clinical trials (Bykov, Eriksson, Bianchi, & Wiman, 2018). Progress has also been made to reactivate nonsense-mutant *TP53.* For example, induced translational readthrough of *TP*53^R213X^ has been shown to reverse the oncogenic effect of lowering *α* due to a truncating mutation (Bidou, Bugaud, Belakhov, Baasov, & Namy, 2017; Floquet, Deforges, Rousset, & Bidou, 2011). Our analysis suggests that concepts like “stabilizing” and “unleashing” p53 must be interpreted in terms of p53’s oscillatory competence, not just its steady-state levels. In this regard, our unified framework of diverse oncogenic alterations in the p53 pathway could guide future therapeutic development to restore p53 oscillations.

Apart from the four parameters considered above, we observed that the remaining parameters (*γ, κ, ϕ* and *ϵ*) could also modulate the dynamic behavior of the p53-Mdm2 system (Fig S3). Although these parameters are not affected by oncogenic alterations, they are potentially modifiable by pharmacological interventions.

## 4 Methods

### Mathematical modeling

Analytic calculations and numerical simulations of the mathematical model were performed in MATLAB (MathWorks). The parameter values we used are shown in Table S1. The justification for these values is as follows: *α, β, γ*: taken from Tsabar et al. (2020); *σ*: taken from Mönke et al. (2017), rescaled to agree with Tsabar et al. (2020); κ: taken from Monke et al. (2017); the parameters *σ, ϵ, ϕ* and *δ* form a group, all affecting Mdm2 dynamics. As a reality check, we verified that the chosen set of parameters produced oscillations in p53 protein levels with the characteristic period of 5-6 hours Lahav et al. (2004), as well as oscillations in *MDM2* transcript and Mdm2 protein levels that are consistent with experimental observations (Hanson et al., 2019). We tested three different sets of parameter values and found that the qualitative behavior of the model was robust under all three sets of parameter values (Table S2; Fig. S4).

### Hopf bifurcation analysis

In dynamical systems theory, a qualitative change in system behavior as a parameter varies is called a bifurcation, and the emergence of a stable limit cycle attractor from a stable equilibrium point as parameters vary is called Poincare-Andronov-Hopf bifurcation, usually shortened to *Hopf bifurcation* (Garfinkel, Guo, & Shevtsov, 2017).

The mathematical signature of Hopf bifurcation is given by linear stability analysis of the equilibrium point(s) of a system, that is, by the eigenvalue structure of the equilibrium point. The Hopf Bifurcation Theorem tells us that when a pair of complex conjugate eigenvalues change from negative real part to positive real part, the equilibrium point will become unstable, and a new stable oscillation, a limit cycle attractor, will be created. Therefore, we can analytically calculate the critical regions for oscillation in parameter space by calculating how the eigenvalue structure depends on those parameters. We used these tools of dynamical systems theory to evaluate the Hopf bifurcation behavior of the p53-Mdm2 model, and to derive the parametric conditions under which p53 oscillations can be sustained.

#### Location of Equilibrium Points

First, we looked for equilibrium points of the p53-Mdm2 system at which p53 (*X_EP_*), *MDM2* transcript (*R_EP_*) and Mdm2 (*Y_EP_*) levels are all positive.

At an equilibrium point, we have 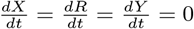. That is,

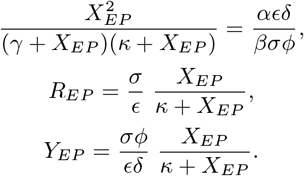

Hence, the equilibrium solutions are:

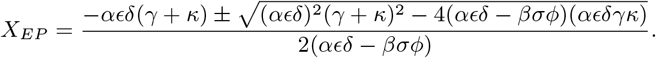

This equation gives an analytic solution of the steady state p53 concentration as a function of the eight model parameters.

#### Stability of Equilibria

Next, we performed linear stability analysis of the system at the equilibrium point in order to derive conditions for the oscillations in the p53-Mdm2 system. The Jacobian matrix evaluated at the equilibrium point has the form:

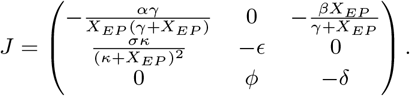

The eigenvalues (*λ*) of the Jacobian matrix are roots to the characteristic (cubic) equation:

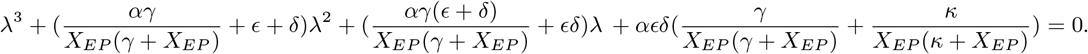

A cubic equation must have at least one real-valued solution. If we denote the real-valued solution as *λ_real_*, we can factor the cubic equation as the following:

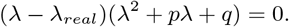

Then the other two roots are: 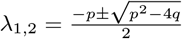. By expanding the factored cubic equation, we derive:

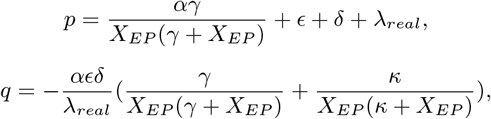

where both *X_EP_* and *λ_real_* are functions of the eight model parameters.

At a Hopf bifurcation point, a pair of complex conjugate eigenvalues passes from having a negative real part to having a positive real part. It follows that a limit cycle attractor is created when the following two conditions are satisfied:

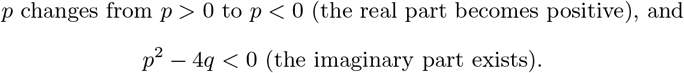

Further details are provided in *SI Appendix*.

All data and code are available at: https://github.com/LingyunXiong/p53_oscillations.

## Author roles

Conceptualization: L.X. and A.G.; Methodology: L.X. and A.G.; Investigation: L.X.; Formal Analysis: L.X. and A.G.; Visualization: L.X.; Writing: L.X. and A.G.; Resources: L.X. and A.G.; Supervision: A.G.

## References

Alexandrova, E. M., Yallowitz, A. R., Li, D., Xu, S., Schulz, R., Proia, D. A., … Moll, U. M. (2015, July). Improving survival by exploiting tumour dependence on stabilized mutant p53 for treatment. Nature, 523(7560), 352–356. Retrieved 2020-11-12, from http://www.nature.com/articles/nature14430 DOI: 10.1038/nature14430

Banin, S., Moyal, L., Shieh, S., Taya, Y., Anderson, C. W., Chessa, L., … Ziv, Y. (1998, September). Enhanced phosphorylation of p53 by ATM in response to DNA damage. Science (New York, N.Y.), 281 (5383), 1674–1677. DOI: 10.1126/science.281.5383.1674

Batchelor, E., Loewer, A., Mock, C., & Lahav, G. (2011, January). Stimulus-dependent dynamics of p53 in single cells. Molecular S’ystems Biology, 7(1), 488. Retrieved 2020-11-06, from https://onlinelibrary.wiley.com/doi/abs/10.1038/msb.2011.20 DOI: 10.1038/msb.2011.20

Batchelor, E., Mock, C. S., Bhan, I., Loewer, A., & Lahav, G. (2008, May). Recurrent Initiation: A Mechanism for Triggering p53 Pulses in Response to DNA Damage. Molecular Cell, 30(3), 277–289. Retrieved 2020-06-11, from https://linkinghub.elsevier.com/retrieve/pii/S1097276508002402 DOI: 10.1016/j.molcel.2008.03.016

Bidou, L., Bugaud, O., Belakhov, V., Baasov, T., & Namy, O. (2017). Characterization of new-generation aminoglycoside promoting premature termination codon readthrough in cancer cells. RNA biology, 14 (3), 378–388. DOI: 10.1080/15476286.2017.1285480

Boiteux, A., Hess, B., & Sel’Kov, E. E. (1980). Creative Functions of Instability and Oscillations in Metabolic Systems. In Current Topics in Cellular Regulation (Vol. 17, pp. 171–203). Elsevier. Retrieved 2020-11-17, from https://linkinghub.elsevier.com/retrieve/pii/B9780121528171500109 DOI: 10.1016/B978-0-12-152817-1.50010-9

Bond, G. L., Hu, W., Bond, E. E., Robins, H., Lutzker, S. G., Arva, N. C., … Levine, A. J. (2004, November). A single nucleotide polymorphism in the MDM2 promoter attenuates the p53 tumor suppressor pathway and accelerates tumor formation in humans. Cell, 119(5), 591–602. DOI: 10.1016/j.cell.2004.11.022

Bougeard, G., Renaux-Petel, M., Flaman, J.-M., Charbonnier, C., Fermey, P., Belotti, M., … Frebourg, T. (2015, July). Revisiting Li-Fraumeni Syndrome From *TP53* Mutation Carriers. Journal of Clinical Oncology, 33(21), 2345–2352. Retrieved 2020-11-25, from https://ascopubs.org/doi/10.1200/JCO.2014.59.5728 DOI: 10.1200/JCO.2014.59.5728

Buhr, E. D., Yoo, S.-H., & Takahashi, J. S. (2010, October). Temperature as a Universal Resetting Cue for Mammalian Circadian Oscillators. Science., 330(6002), 379–385. Retrieved 2020-11-17, from https://www.sciencemag.org/lookup/doi/10.1126/science.1195262 DOI: 10.1126/science.1195262

Bykov, V. J. N., Eriksson, S. E., Bianchi, J., & Wiman, K. G. (2018, February). Targeting mutant p53 for efficient cancer therapy. Nature Reviews Cancer, 18(2), 89–102. Retrieved 2020-11-15, from http://www.nature.com/articles/nrc.2017.109 DOI: 10.1038/nrc.2017.109

Bykov, V. J. N., Issaeva, N., Shilov, A., Hultcrantz, M., Pugacheva, E., Chumakov, P., … Selivanova, G. (2002, March). Restoration of the tumor suppressor function to mutant p53 by a low-molecular-weight compound. Nature Medicine, 8(3), 282–288. DOI: 10.1038/nm0302-282

Chen, D., Shan, J., Zhu, W.-G., Qin, J., & Gu, W. (2010, March). Transcription-independent ARF regulation in oncogenic stress-mediated p53 responses. Nature, 464 (7288), 624–627. Retrieved 2020-11-14, from http://www.nature.com/articles/nature08820 DOI: 10.1038/nature08820

Chen, P., Chen, Y., Bookstein, R., & Lee, W. (1990, December). Genetic mechanisms of tumor suppression by the human p53 gene. Science, 250(4987), 1576–1580. Retrieved 2020-11-17, from https://www.sciencemag.org/lookup/doi/10.1126/science.2274789 DOI: 10.1126/science.2274789

Cheng, Q., Chen, L., Li, Z., Lane, W. S., & Chen, J. (2009, December). ATM activates p53 by regulating MDM2 oligomerization and E3 processivity. The EMBO journal, 28 (24), 3857–3867. DOI: 10.1038/emboj.2009.294

Ciliberto, A., Novak, B., & Tyson, J. J. (2005, March). Steady states and oscillations in the p53/Mdm2 network. Cell Cycle (Georgetown, Tex.), 4 (3), 488–493. DOI: 10.4161/cc.4.3.1548

Donehower, L. A., Soussi, T., Korkut, A., Liu, Y., Schultz, A., Cardenas, M., … Wheeler, D. A. (2019, July). Integrated Analysis of TP53 Gene and Pathway Alterations in The Cancer Genome Atlas. Cell Reports, 28(5), 1370–1384.e5. Retrieved 2020-11-11, from https://linkinghub.elsevier.com/retrieve/pii/S221112471930885X DOI: 10.1016/j.celrep.2019.07.001

Efeyan, A., Garcia-Cao, I., Herranz, D., Velasco-Miguel, S., & Serrano, M. (2006, September). Policing of oncogene activity by p53. Nature, 443(7108), 159–159. Retrieved 2020-11-15, from http://www.nature.com/articles/443159a DOI: 10.1038/443159a

Ferrell, J., Tsai, T.-C., & Yang, Q. (2011, March). Modeling the Cell Cycle: Why Do Certain Circuits Oscillate? Cell, 144 (6), 874–885. Retrieved 2020-11-17, from https://linkinghub.elsevier.com/retrieve/pii/S0092867411002431 DOI: 10.1016/j.cell.2011.03.006

Finlay, C. A., Hinds, P. W., & Levine, A. J. (1989, June). The p53 proto-oncogene can act as a suppressor of transformation. Cell, 57(7), 1083–1093. DOI: 10.1016/0092-8674(89)90045-7

Floquet, C., Deforges, J., Rousset, J.-P., & Bidou, L. (2011, April). Rescue of non-sense mutated p53 tumor suppressor gene by aminoglycosides. Nucleic Acids Research, 39(8), 3350–3362. DOI: 10.1093/nar/gkq1277

Garfinkel, A., Guo, Y., & Shevtsov, J. (2017). Modeling Life: The Mathematics of Biological Systems (1st ed. 2017 ed.). Cham: Springer International Publishing : Imprint: Springer. DOI: 10.1007/978-3319-59731-7

Geva-Zatorsky, N., Rosenfeld, N., Itzkovitz, S., Milo, R., Sigal, A., Dekel, E., … Alon, U. (2006, January). Oscillations and variability in the p53 system. Molecular S’ystems Biology, 2(1). Retrieved 2020-06-11, from https://onlinelibrary.wiley.com/doi/abs/10.1038/msb4100068 DOI: 10.1038/msb4100068

Giacomelli, A. O., Yang, X., Lintner, R. E., McFarland, J. M., Duby, M., Kim, J., … Hahn, W. C. (2018, October). Mutational processes shape the landscape of TP53 mutations in human cancer. Nature Genetics, 50(10), 1381–1387. Retrieved 2020-11-15, from http://www.nature.com/articles/s41588-018-0204-y DOI: 10.1038/s41588-018-0204-y

Glass, L. (2001, March). Synchronization and rhythmic processes in physiology. Nature, 410(6825), 277–284. DOI: 10.1038/35065745

Gu, J., Kawai, H., Nie, L., Kitao, H., Wiederschain, D., Jochemsen, A. G., … Yuan, Z.-M. (2002, May). Mutual dependence of MDM2 and MDMX in their functional inactivation of p53. The Journal of Biological Chemistry, 277(22), 19251–19254. DOI: 10.1074/jbc.C200150200

Hanson, R. L., Porter, J. R., & Batchelor, E. (2019, April). Protein stability of p53 targets determines their temporal expression dynamics in response to p53 pulsing. Journal of Cell Biology, 218 (4), 1282–1297. Retrieved 2020-11-21, from https://rupress.org/jcb/article/218/4/1282/61849/Protein-stability-of-p53-targets-determines-their DOI:10.1083/jcb.201803063

Haupt, Y., Maya, R., Kazaz, A., & Oren, M. (1997, May). Mdm2 promotes the rapid degradation of p53. Nature, 387(6630), 296–299. DOI: 10.1038/387296a0

Heltberg, M. L., Chen, S.-h., Jiménez, A., Jambhekar, A., Jensen, M. H., & Lahav, G. (2019, December). Inferring Leading Interactions in the p53/Mdm2/Mdmx Circuit through Live-Cell Imaging and Modeling. Cell Systems, 9(6), 548–558.e5. Retrieved 2020-08-15, from https://linkinghub.elsevier.com/retrieve/pii/S2405471219303862 DOI: 10.1016/j.cels.2019.10.010

Hu, W., Feng, Z., Ma, L., Wagner, J., Rice, J. J., Stolovitzky, G., & Levine, A. J. (2007, March). A single nucleotide polymorphism in the MDM2 gene disrupts the oscillation of p53 and MDM2 levels in cells. Cancer Research, 67(6), 2757–2765. DOI: 10.1158/0008-5472.CAN-06-2656

Kamijo, T., Bodner, S., van de Kamp, E., Randle, D. H., & Sherr, C. J. (1999, May). Tumor spectrum in ARF-deficient mice. Cancer Research, 59(9), 2217–2222.

Kamijo, T., Zindy, F., Roussel, M. F., Quelle, D. E., Downing, J. R., Ashmun, R. A., … Sherr, C. J. (1997, November). Tumor Suppression at the Mouse INK4a Locus Mediated by the Alternative Reading Frame Product p19 ARF. Cell, 91 (5), 649–659. Retrieved 2020-11-17, from https://linkinghub.elsevier.com/retrieve/pii/S0092867400804523 DOI: 10.1016/S0092-8674(00)80452-3

Kasper, E., Angot, E., Colasse, E., Nicol, L., Sabourin, J.-C., Adriouch, S., … Bougeard, G. (2018, September). Contribution of genotoxic anticancer treatments to the development of multiple primary tumours in the context of germline TP53 mutations. European Journal of Cancer, 101, 254–262. Retrieved 2020-11-25, from https://linkinghub.elsevier.com/retrieve/pii/S0959804918309031 DOI: 10.1016/j.ejca.2018.06.011

Kato, S., Han, S.-Y., Liu, W., Otsuka, K., Shibata, H., Kanamaru, R., & Ishioka, C. (2003, July). Understanding the function-structure and function-mutation relationships of p53 tumor suppressor protein by high-resolution missense mutation analysis. Proceedings of the National Academy of Sciences of the United States of America, 100(14), 8424–8429. DOI: 10.1073/pnas.1431692100

Khosravi, R., Maya, R., Gottlieb, T., Oren, M., Shiloh, Y., & Shkedy, D. (1999, December). Rapid ATM-dependent phosphorylation of MDM2 precedes p53 accumulation in response to DNA damage. Proceedings of the National Academy of Sciences of the United States of America, 96(26), 14973–14977. DOI: 10.1073/pnas.96.26.14973

Knijnenburg, T. A., Wang, L., Zimmermann, M. T., Chambwe, N., Gao, G. F., Cherniack, A. D., … Mariamidze, A. (2018, April). Genomic and Molecular Landscape of DNA Damage Repair Deficiency across The Cancer Genome Atlas. Cell Reports, 23(1), 239–254.e6. Retrieved 2020-11-11, from https://linkinghub.elsevier.com/retrieve/pii/S2211124718304376 DOI: 10.1016/j.celrep.2018.03.076

Kotler, E., Shani, O., Goldfeld, G., Lotan-Pompan, M., Tarcic, O., Gershoni, A., … Segal, E. (2018, July). A Systematic p53 Mutation Library Links Differential Functional Impact to Cancer Mutation Pattern and Evolutionary Conservation. Molecular Cell, 71 (1), 178–190.e8. Retrieved 2020-11-15, from https://linkinghub.elsevier.com/retrieve/pii/S1097276518304544 DOI: 10.1016/j.molcel.2018.06.012

Lahav, G., Rosenfeld, N., Sigal, A., Geva-Zatorsky, N., Levine, A. J., Elowitz, M. B., & Alon, U. (2004, February). Dynamics of the p53-Mdm2 feedback loop in individual cells. Nature Genetics, 36(2), 147–150. Retrieved 2020-08-15, from http://www.nature.com/articles/ng1293 DOI: 10.1038/ng1293

Lambert, J. M. R., Gorzov, P., Veprintsev, D. B., Söderqvist, M., Segerbäck, D., Bergman, J., … Bykov, V. J. N. (2009, May). PRIMA-1 reactivates mutant p53 by covalent binding to the core domain. Cancer Cell, 15(5), 376–388. DOI: 10.1016/j.ccr.2009.03.003

Lan, K.-H., Sheu, M.-L., Hwang, S.-J., Yen, S.-H., Chen, S.-Y., Wu, J.-C., … Lee, S.-D. (2002, July). HCV NS5A interacts with p53 and inhibits p53-mediated apoptosis. Oncogene, 21 (31), 4801–4811. Retrieved 2020-10-30, from http://www.nature.com/articles/1205589 DOI: 10.1038/sj.onc.1205589

Lane, D. P., & Crawford, L. V. (1979, March). T antigen is bound to a host protein in SV40-transformed cells. Nature, 278(5701), 261–263. DOI: 10.1038/278261a0

Lev Bar-Or, R., Maya, R., Segel, L. A., Alon, U., Levine, A. J., & Oren, M. (2000, October). Generation of oscillations by the p53-Mdm2 feedback loop: A theoretical and experimental study. Proceedings of the National Academy of Sciences, 97(21), 11250–11255. Retrieved 2020-11-25, from http://www.pnas.org/cgi/doi/10.1073/pnas.210171597 DOI: 10.1073/pnas.210171597

Levine, A. J. (1997, February). p53, the cellular gatekeeper for growth and division. Cell, 88(3), 323–331. DOI: 10.1016/s0092-8674(00)81871-1

Levine, J. H., Lin, Y., & Elowitz, M. B. (2013, December). Functional Roles of Pulsing in Genetic Circuits. Science, 342(6163), 1193–1200. Retrieved 2020-11-21, from https://www.sciencemag.org/lookup/doi/10.1126/science.1239999 DOI: 10.1126/science.1239999

Li, D., Marchenko, N. D., & Moll, U. M. (2011, December). SAHA shows preferential cytotoxicity in mutant p53 cancer cells by destabilizing mutant p53 through inhibition of the HDAC6-Hsp90 chaperone axis. Cell Death & Differentiation, 18(12), 1904–1913. Retrieved 2020-11-12, from http://www.nature.com/articles/cdd201171 DOI: 10.1038/cdd.2011.71

Li, D., Marchenko, N. D., Schulz, R., Fischer, V., Velasco-Hernandez, T., Talos, F., & Moll, U. M. (2011, May). Functional Inactivation of Endogenous MDM2 and CHIP by HSP90 Causes Aberrant Stabilization of Mutant p53 in Human Cancer Cells. Molecular Cancer Research, 9(5), 577–588. Retrieved 2020-11-11, from http://mcr.aacrjournals.org/cgi/doi/10.1158/1541-7786.MCR-10-0534 DOI: 10.1158/1541-7786.MCR-10-0534

Linzer, D. I., & Levine, A. J. (1979, May). Characterization of a 54K dalton cellular SV40 tumor antigen present in SV40-transformed cells and uninfected embryonal carcinoma cells. Cell, 17(1), 43–52. DOI: 10.1016/0092-8674(79)90293-9

Livingstone, L. R., White, A., Sprouse, J., Livanos, E., Jacks, T., & Tlsty, T. D. (1992, September). Altered cell cycle arrest and gene amplification potential accompany loss of wild-type p53. Cell, 70(6), 923–935. Retrieved 2020-11-11, from https://linkinghub.elsevier.com/retrieve/pii/0092867492902436 DOI: 10.1016/0092-8674(92)90243-6

Ma, L., Wagner, J., Rice, J. J., Hu, W., Levine, A. J., & Stolovitzky, G. A. (2005, October). A plausible model for the digital response of p53 to DNA damage. Proceedings of the National Academy of Sciences, 102(40), 14266–14271. Retrieved 2020-11-20, from http://www.pnas.org/cgi/doi/10.1073/pnas.0501352102 DOI: 10.1073/pnas.0501352102

Macedo, G. S., Araujo Vieira, I., Brandalize, A. P., Giacomazzi, J., Inez Palmero, E., Volc, S., … Ashton-Prolla, P. (2016, March). Rare germline variant (rs78378222) in the TP53 3’ UTR: Evidence for a new mechanism of cancer predisposition in Li-Fraumeni syndrome. Cancer Genetics, 209(3), 97–106. Retrieved 2020-10-12, from https://linkinghub.elsevier.com/retrieve/pii/S221077621600003X DOI: 10.1016/j.cancergen.2015.12.012

Maya, R., Balass, M., Kim, S. T., Shkedy, D., Leal, J. F., Shifman, O., … Oren, M. (2001, May). ATM-dependent phosphorylation of Mdm2 on serine 395: role in p53 activation by DNA damage. Genes & Development, 15(9), 1067–1077. DOI: 10.1101/gad.886901

Momand, J., Zambetti, G. P., Olson, D. C., George, D., & Levine, A. J. (1992, June). The mdm-2 oncogene product forms a complex with the p53 protein and inhibits p53-mediated transactivation. Cell, 69(7), 1237–1245. Retrieved 2020-11-11, from https://linkinghub.elsevier.com/retrieve/pii/009286749290644R DOI: 10.1016/0092-8674(92)90644-R

Mönke, G., Cristiano, E., Finzel, A., Friedrich, D., Herzel, H., Falcke, M., & Loewer, A. (2017, June). Excitability in the p53 network mediates robust signaling with tunable activation thresholds in single cells. Scientific Reports, 7(1), 46571. Retrieved 2020-11-15, from http://www.nature.com/articles/srep46571 DOI: 10.1038/srep46571

Noble, D. (2006). The music of life: biology beyond the genome. Oxford; New York: Oxford University Press. (OCLC: ocm64554661)

Park, D. J., Nakamura, H., Chumakov, A. M., Said, J. W., Miller, C. W., Chen, D. L., & Koeffler, H. P. (1994, July). Transactivational and DNA binding abilities of endogenous p53 in p53 mutant cell lines. Oncogene, 9(7), 1899–1906.

Pei, D., Zhang, Y., & Zheng, J. (2012, March). Regulation of p53: a collaboration between Mdm2 and Mdmx. Oncotarget, 3(3), 228–235. DOI: 10.18632/oncotarget.443

Peng, Y., Chen, L., Li, C., Lu, W., & Chen, J. (2001, November). Inhibition of MDM2 by hsp90 contributes to mutant p53 stabilization. The Journal of Biological Chemistry, 276(44), 40583–40590. DOI: 10.1074/jbc.M102817200

Pise-Masison, C. A., Choi, K.-S., Radonovich, M., Dittmer, J., Kim, S.-J., & Brady, J. N. (1998, February). Inhibition of p53 Transactivation Function by the Human T-Cell Lymphotropic Virus Type 1 Tax Protein. Journal of Virology, 72(2), 1165–1170. Retrieved 2020-10-30, from https://JVI.asm.org/content/72/2/1165 DOI: 10.1128/JVI.72.2.1165-1170.1998

Pomerantz, J., Schreiber-Agus, N., Liégeois, N. J., Silverman, A., Alland, L., Chin, L., … DePinho, R. A. (1998, March). The Ink4a Tumor Suppressor Gene Product, p19Arf, Interacts with MDM2 and Neutralizes MDM2’s Inhibition of p53. Cell, 92(6), 713–723. Retrieved 2020-11-11, from https://linkinghub.elsevier.com/retrieve/pii/S0092867400814002 DOI: 10.1016/S0092-8674(00)81400-2

Post, S. M., Quintá-Cardama, A., Pant, V., Iwakuma, T., Hamir, A., Jackson, J. G., … Lozano, G. (2010, September). A High-Frequency Regulatory Polymorphism in the p53 Pathway Accelerates Tumor Development. Cancer Cell, 18(3), 220–230. Retrieved 2020-11-06, from https://linkinghub.elsevier.com/retrieve/pii/S153561081000303X DOI: 10.1016/j.ccr.2010.07.010

Purvis, J. E., Karhohs, K. W., Mock, C., Batchelor, E., Loewer, A., & Lahav, G. (2012, June). p53 Dynamics Control Cell Fate. Science, 336(6087), 1440–1444. Retrieved 2020-06-11, from https://www.sciencemag.org/lookup/doi/10.1126/science.1218351 DOI: 10.1126/science.1218351

Salim, K. Y., Maleki Vareki, S., Danter, W. R., & Koropatnick, J. (2016, July). COTI-2, a novel small molecule that is active against multiple human cancer cell lines in vitro and in vivo. Oncotarget, 7(27), 41363–41379. DOI: 10.18632/oncotarget.9133

Sarnow, P., Ho, Y. S., Williams, J., & Levine, A. J. (1982, February). Adenovirus E1b-58kd tumor antigen and SV40 large tumor antigen are physically associated with the same 54 kd cellular protein in transformed cells. Cell, 28(2), 387–394. DOI: 10.1016/0092-8674(82)90356-7

Scheffner, M., Werness, B. A., Huibregtse, J. M., Levine, A. J., & Howley, P. M. (1990, December). The E6 oncoprotein encoded by human papillomavirus types 16 and 18 promotes the degradation of p53. Cell, 63(6), 1129–1136. DOI: 10.1016/0092-8674(90)90409-8

Sherr, C. J. (1998, October). Tumor surveillance via the ARF-p53 pathway. Genes & Development, 12(19), 2984–2991. Retrieved 2020-11-11, from http://www.genesdev.org/cgi/doi/10.1101/gad.12.19.2984 DOI: 10.1101/gad.12.19.2984

Shieh, S. Y., Ikeda, M., Taya, Y., & Prives, C. (1997, October). DNA damage-induced phosphorylation of p53 alleviates inhibition by MDM2. Cell, 91 (3), 325–334. DOI: 10.1016/s0092-8674(00)80416-x

Stewart-Ornstein, J., & Lahav, G. (2017, April). p53 dynamics in response to DNA damage vary across cell lines and are shaped by efficiency of DNA repair and activity of the kinase ATM. Science Signaling, 10(476). DOI: 10.1126/scisignal.aah6671

Stommel, J. M., & Wahl, G. M. (2004, April). Accelerated MDM2 auto-degradation induced by DNA-damage kinases is required for p53 activation. The EMBO journal, 23(7), 1547–1556. DOI: 10.1038/sj.emboj.7600145

Tsabar, M., Mock, C. S., Venkatachalam, V., Reyes, J., Karhohs, K. W., Oliver, T. G., … Lahav, G. (2020, August). A Switch in p53 Dynamics Marks Cells That Escape from DSB-Induced Cell Cycle Arrest. Cell Reports, 32(5), 107995. Retrieved 2020-08-15, from https://linkinghub.elsevier.com/retrieve/pii/S2211124720309803 DOI: 10.1016/j.celrep.2020.107995

Tu, B. P., Kudlicki, A., Rowicka, M., & McKnight, S. L. (2005, November). Logic of the yeast metabolic cycle: temporal compartmentalization of cellular processes. Science (New York, N.Y.), 310(5751), 1152–1158. DOI: 10.1126/science.1120499

Venkatachalam, S. (1998, August). Retention of wild-type p53 in tumors from p53 heterozygous mice: reduction of p53 dosage can promote cancer formation. The EMBO Journal, 17(16), 4657–4667. Retrieved 2020-11-11, from http://emboj.embopress.org/cgi/doi/10.1093/emboj/17.16.4657 DOI: 10.1093/emboj/17.16.4657

Wagner, J., Ma, L., Rice, J. J., Hu, W., Levine, A. J., & Stolovitzky, G. A. (2005, September). p53-Mdm2 loop controlled by a balance of its feedback strength and effective dampening using ATM and delayed feedback. Systems Biology, 152(3), 109–118. DOI: 10.1049/ip-syb:20050025

Wang, X., & Sun, Q. (2017, January). TP53 mutations, expression and interaction networks in human cancers. Oncotarget, 8(1), 624–643. DOI: 10.18632/oncotarget.13483

Wang, X. W., Forrester, K., Yeh, H., Feitelson, M. A., Gu, J. R., & Harris, C. C. (1994, March).Hepatitis B virus X protein inhibits p53 sequence-specific DNA binding, transcriptional activity, and association with transcription factor ERCC3. Proceedings of the National Academy of Sciences of the United States of America, 91 (6), 2230–2234. DOI: 10.1073/pnas.91.6.2230

Weber, J. D., Taylor, L. J., Roussel, M. F., Sherr, C. J., & Bar-Sagi, D. (1999, May). Nucleolar Arf sequesters Mdm2 and activates p53. Nature Cell Biology, 1 (1), 20–26. Retrieved 2020-11-13, from http://www.nature.com/articles/ncb059W20 DOI: 10.1038/8991

Werness, B. A., Levine, A. J., & Howley, P. M. (1990, April). Association of human papillomavirus types 16 and 18 E6 proteins with p53. Science (New York, N.Y.), 2f8(4951), 76–79. DOI: 10.1126/science.2157286

Yi, F., Saha, A., Murakami, M., Kumar, P., Knight, J. S., Cai, Q., … Robertson, E. S. (2009, June). Epstein-Barr virus nuclear antigen 3C targets p53 and modulates its transcriptional and apoptotic activities. Virology, 388(2), 236–247. DOI: 10.1016/j.virol.2009.03.027

Zhang, P., Kitchen-Smith, I., Xiong, L., Stracquadanio, G., Brown, K., Richter, P., … Bond, G. (2019, November). *Germline and somatic genetic variants in the p53 pathway interact to affect cancer risk, progression and drug response* (preprint). Cancer Biology. Retrieved 2020-06-11, from http://biorxiv.org/lookup/doi/10.1101/835918 DOI: 10.1101/835918

Zhang, Y., Xiong, Y., & Yarbrough, W. G. (1998, March). ARF Promotes MDM2 Degradation and Stabilizes p53: ARF-INK4a Locus Deletion Impairs Both the Rb and p53 Tumor Suppression Pathways. Cell, 92(6), 725–734. Retrieved 2020-11-11, from https://linkinghub.elsevier.com/retrieve/pii/S0092867400814014 DOI: 10.1016/S0092-8674(00)81401-4

